# The Primate-Specific Presence of Interferon Regulatory Factor-5 Pseudogene 1

**DOI:** 10.1101/2024.02.07.579370

**Authors:** Avery Marquis, Vanessa Hubing, Chanasei Ziemann, Etsuko N. Moriyama, Luwen Zhang

## Abstract

Interferon regulatory factor 5 (IRF5) is a critical transcription factor, regulating the production of inflammatory cytokines and interferons. Dysregulation of IRF5 has been linked to various autoimmune and inflammatory diseases. Pseudogenes may exhibit gene regulatory functions in various mechanisms. In this study, we find the human IRF5 pseudogene 1 (IRF5P1) is a chimeric processed pseudogene containing sequences derived from multiple sources. Remarkably, IRF5P1 is specific to higher primates such as apes and humans. We propose that IRF5P1 may have originated through the ancient integration of a retroviral sequence containing IRF5 mRNA from other animals. Interestingly, IRF5P1 overlaps with another gene, triple QxxK/R motif containing (TRIQK), and the antisense strand of IRF5P1 is predominantly expressed, likely as part of the TRIQK pre-mRNA. The antisense IRF5P1 RNA may regulate IRF5 expression through complementary binding to the IRF5 mRNA, and variants in the IRF5 gene may modulate this interaction. Our analysis suggests that IRF5P1 likely had positive effects on primate evolution, making IRF5P1 a favored component to maintain in the primate lineage, and rendering the pseudogene primate specific. Further studies are essential to elucidate the potential for IRF5P1 to participate in regulatory networks governing IRF5 activity and inflammation and innate immunity.

## Introduction

Interferon regulatory factor 5 (IRF5) is a transcription factor that plays a key role in the regulation of inflammatory and immune responses. IRF5 is a member of the IRF family of transcription factors, which contain a conserved DNA-binding domain that binds to interferon-stimulated response elements found in the promoters of interferon target genes (Barnes, Lubyova, & Pitha, 2002; Jefferies, 2019; Tamura, Yanai, Savitsky, & Taniguchi, 2008; Taniguchi, Ogasawara, Takaoka, & Tanaka, 2001; Yanai, Negishi, & Taniguchi, 2012). Upon activation, IRF5 translocates to the nucleus where it regulates expression of downstream target genes involved in type I interferon signaling, cytokine production, cell proliferation, and apoptosis. Dysregulation of IRF5 has been implicated in several autoimmune and inflammatory diseases (Almuttaqi & Udalova, 2019; Jefferies, 2019). The mechanisms by which IRF5 genetic variants contribute to autoimmunity and inflammation continues to be an important avenue of research, as IRF5 may represent a promising therapeutic target for treating these diseases (Almuttaqi & Udalova, 2019).

Pseudogenes are DNA sequences that resemble functional genes but are generally thought to have lost their protein-coding ability or gene expression potential (Tutar, 2012). Pseudogenes are ubiquitous in genomes across taxa and they arise primarily from gene duplication or retrotransposition (Li, Yang, & Wang, 2013; Tutar, 2012). Although initially thought of as nonfunctional genetic elements, recent research has uncovered diverse functional roles for pseudogenes. For example, some retain the ability to produce RNAs that regulate expression levels of functional progenitor genes or even produce truncated proteins with novel activities (Pink et al., 2011; Poliseno et al., 2010). Other pseudogenes play roles in regulating gene expression by acting as decoys for microRNAs that would otherwise suppress mRNAs from related coding genes (Hu, Yang, & Mo, 2018; Poliseno, 2012; Rapicavoli et al., 2013). There is also evidence that pseudogene transcription may elicit autoimmune responses in some diseases (Brooks, 2005; Rapicavoli et al., 2013). Therefore, pseudogenes exhibit functionality ranging from gene expression regulation to generation of immunological epitopes.

We have analyzed the IRF5 pseudogene 1 (IRF5P1) gene in a systemic manner and found that IRF5P1 has high DNA sequence similarity to two mRNA sequences in the functional domains in IRF5. The DNA sequences of IRF5P1 exhibit a mixed origin and likely originated approximately 60 million years ago. Because IRF5P1 antisense RNA are at least transiently expressed, evolutionary conservation in primate genomes implies selection for functional non-coding roles. Studying IRF5P1 may reveal non-coding regulatory functions or novel peptide products relevant to IRF5 biology.

## Results

### IRF5P1 is a chimeric processed pseudogene containing DNA sequences derived from multiple sources

We divided the DNA sequence of IRF5P1 into four sections based on sequence homologies to IRF5 (Figure 1A). Analysis of different regions shows that section 1 (nucleotides 1-98) is apparently present only in some primates (Figure 1B; Supplemental Figure 1) and appears to be primate specific. Section 3 matches fragmented DNA from various other animal species, including cats, dogs, and whales, providing evidence of the integration of diverse external DNA (Figure 1D). Additionally, some IRF5P1-specific sequences are also present (Supplemental Figure 1). Sections 2 and 4 exhibit similarity to IRF5 mRNAs from various species (Figure 1C and 1E). Section 2 also has sequence homology to other animal’ genomic sequences such as whales (Figure 1C). Those genomic sequences are not the exon sequences of the IRF5 genes in the organisms (data not shown). Despite displaying partial sequence homologies to the human IRF5 gene, IRF5P1 harbors multiple out-of-frame and deletion/inserting mutations that hinder its ability to produce a functional protein product. The corresponding hIRF5 amino acid sequences in the sections are shown (Figure 1A). Multiple exons are needed to cover the corresponding Sections 2 and 4 sequences in humans, whales and other mammals (Supplemental Figure 2). In addition, Section 4 has homology to the 3’ untranslated region (UTR) of the IRF5 gene (Figure 1A). The fact that the fragment matches mostly IRF5 mRNA covering multiple exons rather than genomic DNA in the sections 2 and 4 suggests IRF5P1 is specifically a “processed pseudogene” - the introns have been removed, as is typical of mRNAs (Tutar, 2012). All the data suggest that IRF5P1 is a chimeric processed pseudogene with a mixed origin.

**FIGURE 1:**
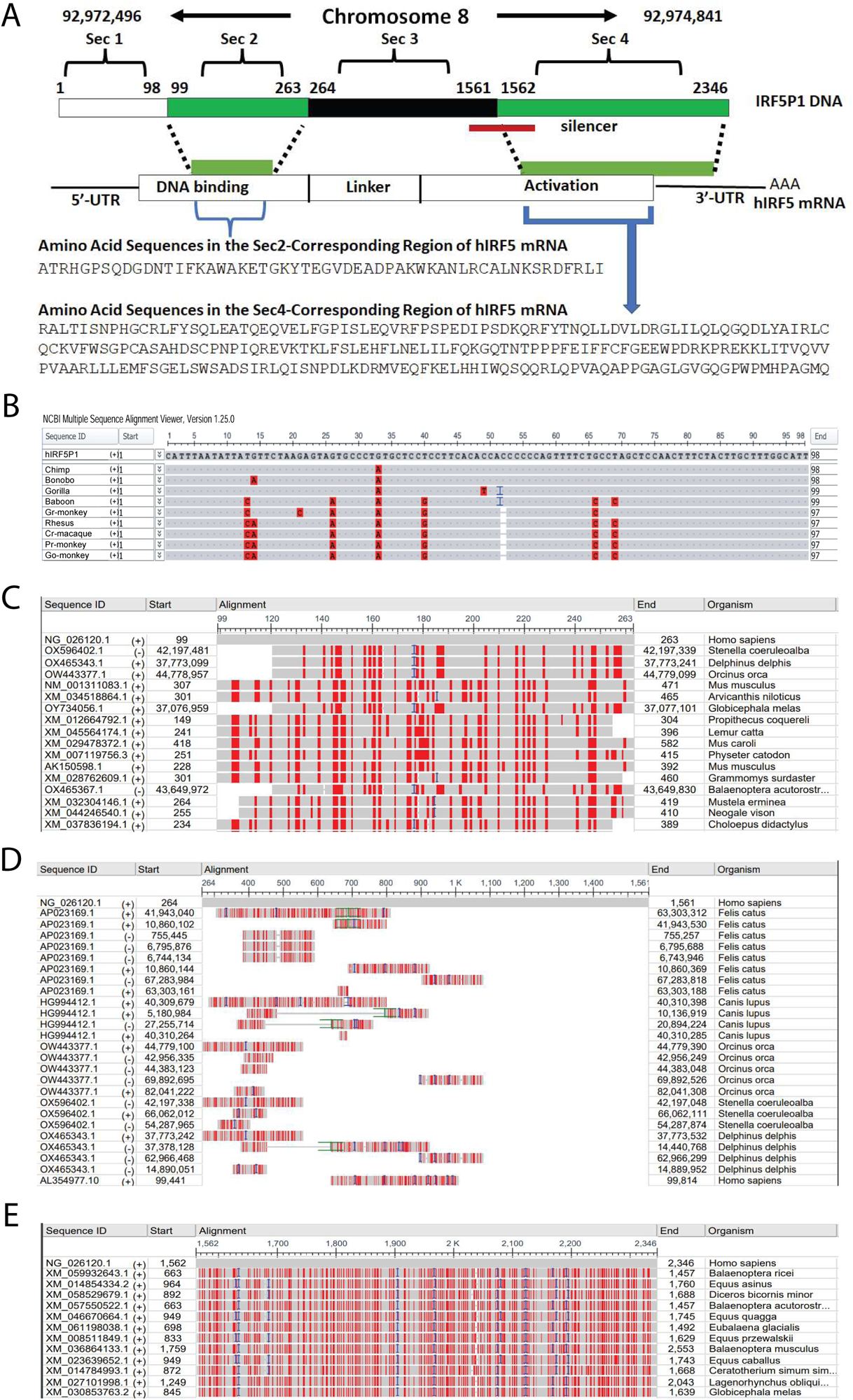
A. Schematic Diagram of IRF5P1 and Sequence Analysis. **Panel A: Diagram of the human IRF5P1 gene on Chromosome 8 (NG_026120.1).** Chromosome number and coordinates are provided, and the IRF5P1 DNA are divided into four sections: Section 1 (Sec 1; 1-98), Section 2 (Sec 2; 99-263), Section 3 (Sec 3; 264-1561), and Section 4 (Sec 4; 1562-2346). A red line below the IRF5P1 DNA represents a silencer. Human IRF5 (hIRF5) mRNA and the corresponding protein coding regions, as well as functional domains, are shown. The amino acid sequences in hIRF5 corresponding to Sec 2 and Sec 4 are displayed. **Panel B: Comparison of hIRF5P1 to Primate Nucleotide Sequences within Section 1 (Sec 1)**. The Sec 1 sequence of IRF5P1 undergoes Blastn analysis, and results are presented using the NCBI Multiple Sequence Alignment Viewer. Similarities and differences are outlined, with sequence IDs corresponding to primate IRF5P1 common names or abbreviations. Cr-macaque: crab-eating macaque; Go-monkey: golden snub-nose monkey; Gr-monkey: green monkey; Pr-monkey: proboscis monkey. Starting and ending sites in human IRF5P1 and other primates are provided. **Panels C-E: Comparison of hIRF5P1 Sections 2-4 with Other Sequences**. Panels C, D, and E focus on comparing hIRF5P1 Section 2 (99-263), Section 3 (264-1561), and Section 4 (1562-2346) with their similar sequences, respectively. Blastn is employed for the sequence search, and the NCBI Multiple Sequence Alignment Viewer aids in comparison and visualization. The table includes accession numbers, start and end coordinates, and organism names. Alignment rows visualize missing (white), homologous (grey), dissimilar (red), and insertion/deletion regions.

### IRF5P1 is a Primate-Specific Pseudogene

While absent in some primates like lemurs and bushbabies, IRF5P1 is present in multiple primate species, including some New World monkeys such as marmosets and is conserved across Old World monkeys as well as apes and humans (Supplemental Table 1 and Supplemental Figure 1). Tarsiers have a gene with great similarity to IRF5P1, but it is missing section 1 (Supplemental Figure 1). However, phylogenetically, the tarsier appears to be an outgroup compared to other IRF5P1s, and rhino has a gene that is closer to other IRF5P1s than the tarsier (Figure 2A). In conclusion, distribution analysis delineates IRF5P1 as a predominantly primate-specific processed pseudogene that originated in the ancient ancestors of extant primates and remained preserved through later primate evolution, including humans.

**FIGURE 2:**
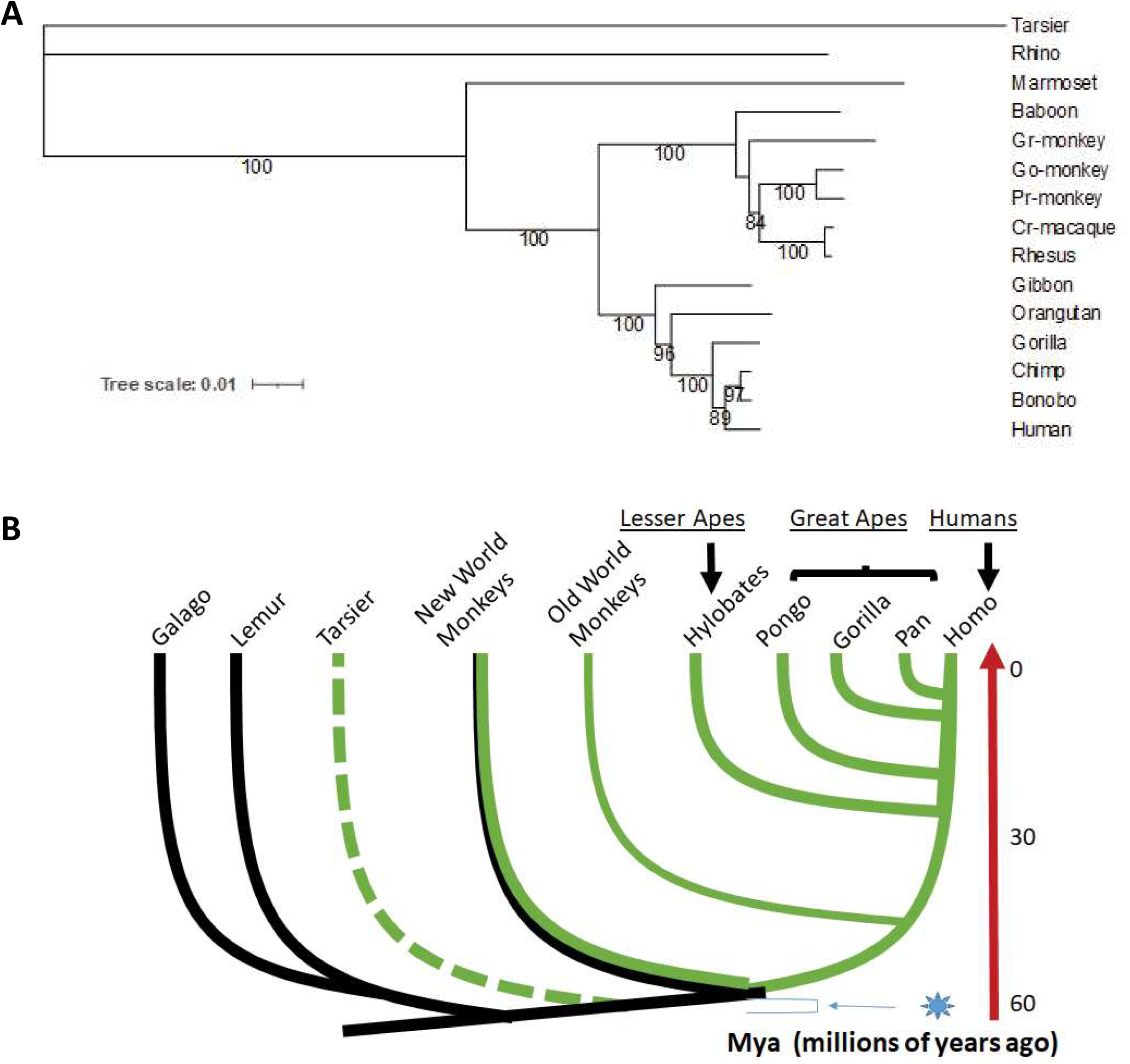
Emergence of IRF5P1 during Evolution. **Panel A**: Phylogenetic relationships among IRF5P1 in various species. The maximum likelihood phylogeny was reconstructed using IRF5P1 nucleotide sequences from 15 species. The Tarsier and Rhino IRF5P1 sequences served as the outgroup. Internal nodes supported by 70% or higher bootstrap values are indicated by numerical values on nodes. Refer to Supplemental Table 1 for the nucleotide sequences used in the study. **Panel B**: The emergence of IRF5P1 during primate evolution. The evolutionary relationship among the primate species in this study is illustrated. Green lines indicate the lineages where the acquisition of IRF5P1 occurred. Not all New World monkeys have IRF5P1, as indicated by the black and green mixed-colored line. The dashed green line leading to the tarsier represents uncertainty regarding the presence of the IRF5P1 gene. Phylogenetic relationships and age estimates for diversifications among primates are approximated based on references (Chapman & Lambert, 2000; Finstermeier et al., 2013; Lucas et al., 2003; Pääbo, 2003; Page, Chiu, & Goodman, 1999; Pecon-Slattery, 2014). Numbers on the right represent approximate divergence times in million years ago (Mya). It is proposed that a virus containing IRF5P1-like sequences infected the ancestors of primates around 60 Mya.

### A Retroviral Event might be a Potential Contributor to the Lineage-Specific Presence of IRF5P1

To study the relationship between hIRF5 and IRF5P1, we analyzed the phylogenetic relationship among mammalian IRF5s concerning IRF5P1. The IRF5 mRNA sequences most closely related to IRF5P1 were determined based on e-values and percentage identities in Blastn search (Figure 1C and 1E; Supplemental Figure 3). The homologous sequences in sections 2 and 4 were studied separately in closely related IRFs. Phylogenetically, the hIRF5 mRNA sequence is an outgroup from other IRF5 mRNA sequences in section 4, and more distantly related to IRF5P1 (Supplemental Figure 3C). In addition, from the available genomic data, IRF5 has extensive splicing events for RNA processing for various animals and both section 2 and 4 sequences are generated through multiple splicing (Supplemental Figure 2). Those data collectively suggest that IRF5P1 was unlikely to be generated by host gene duplication, but a mRNA transcript from foreign IRF5 gene integrated into the genome by reverse transcription at some point. Furthermore based on association among IRF5P1 and IRF5 mRNA sequences from whales, and others (Supplemental Figure 2), it is hypothesized that an ancient retrovirus carrying IRF5 mRNA from the ancestors of those animals infected primate ancestors around 60 million years ago, leading to the horizontal transfer of the gene into the primate lineage (Figure 2B) (Lavialle et al., 2013). IRF5P1 might have once been a functional gene in primates, but it has since lost its function. This loss could have occurred through various mechanisms, such as mutations disrupting the gene’s coding sequences and recombination events.

### Expression of antisense of IRF5P1 RNA in cells

Whether IRF5P1 expression is present was addressed using the existing database. We first noticed that IRF5P1 is in the middle of another gene (Figure 3). The triple QxxK/R motif-containing (TRIQK) is a gene located on human chromosome 8 and overlaps with IRF5P1. The pre-mRNA of TRIQK, but not the mature one, contains the antisense sequence of IRF5P1 (Figure 3A). Indeed, IRF5P1 is expressed in human cells based on RNA expression data (Figure 3B). However, there is minimal RNA expression of the pseudogene in healthy individuals based on another database, and the neighboring TRIQK gene exhibits significant RNA expression (Supplemental Figure 4). Because the data in Supplemental Figure 4 are based on “sense” RNA sequences, these expression data collectively indicate that the expression of the antisense strand of IRF5P1 RNA is predominantly form in human cells (Figure 3B, Supplemental Figure 4). Of note, there is an experimentally verified transcriptional silencer sequence present in the IRF5P1 gene (Figure 1A) (Pang & Snyder, 2020), which may explain the lack of sense IRF5P1 RNA expression in the cells.

**FIGURE 3:**
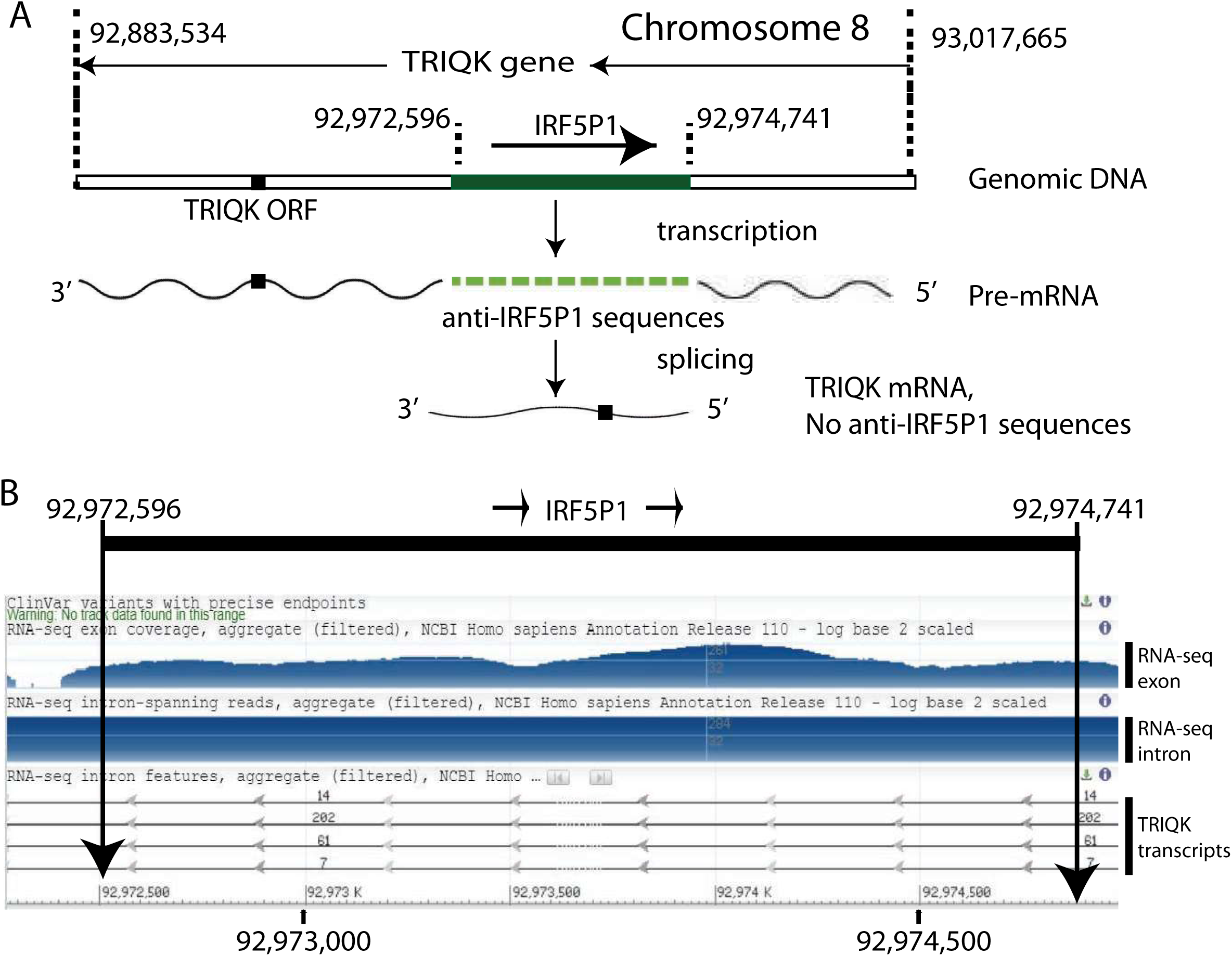
Expression of antisense IRF5P1 in Human Cells. **Panel A: Structure of the TRIQK gene.** Genomic DNA fragment containing TRIQK gene on chromosome 8 at coordinates 93,017,665 and 92,883,534 is as shown. The IRF5P1 gene is also shown. Horizontal lines indicate the directions of the genes. A solid square represents the open reading frame (ORF) of TRIQK. A solid green line represents the IRF5P1 gene. Pre-mRNA and mature mRNA are depicted as waved lines. The dashed green wave line represents the antisense of IRF5P1. The antisense sequence is apparently not present in mature TRIQK mRNA. **Panel B: Transcripts for antisense IRF5P1 in human cells.** Aggregated RNA-seq data for the expression on exon and intron regions are shown. The corresponding nucleotide coordinates are shown, and the IRF5P1 region is indicated. The bottom shows the TRIQK RNA transcripts with indicated direction. The image was captured from the NCBI gene description for IRF5P1 (Gene ID: 100420406, updated on 10-Oct-2023).

### The antisense of IRF5P1 RNA may play a role in regulating IRF5

Because the hIRF5 gene and IRF5P1 have significant homology in nucleotide sequences, it is reasonable to assume that the antisense IRF5P1 RNA regulates IRF5 expression and function through directly binding to IRF5 mRNA. Interestingly, a portion of the 3’ UTR in hIRF5 has significant homology to IRF5P1 (Figure 1A), and we found there are multiple human variants in this region of the DNA (Figure 4). Of note, some variants may directly affect the putative interactions between the IRF5 UTR and IRF5P1 antisense RNA. A C/T variant (rs79645063) may disrupt the potential binding of the IRF5 UTR to IRF5P1 antisense RNA (Figure 4B). Many variants, including deletions and additions (blue bars in Figure 4A) in this UTR region offer various possible ways to regulate IRF5 through antisense RNA-mediated effects. Other homologous regions in sections 2 and 4 may have a similar effect and furthermore human nucleotide variants may affect the amino acid identities.

**FIGURE 4:**
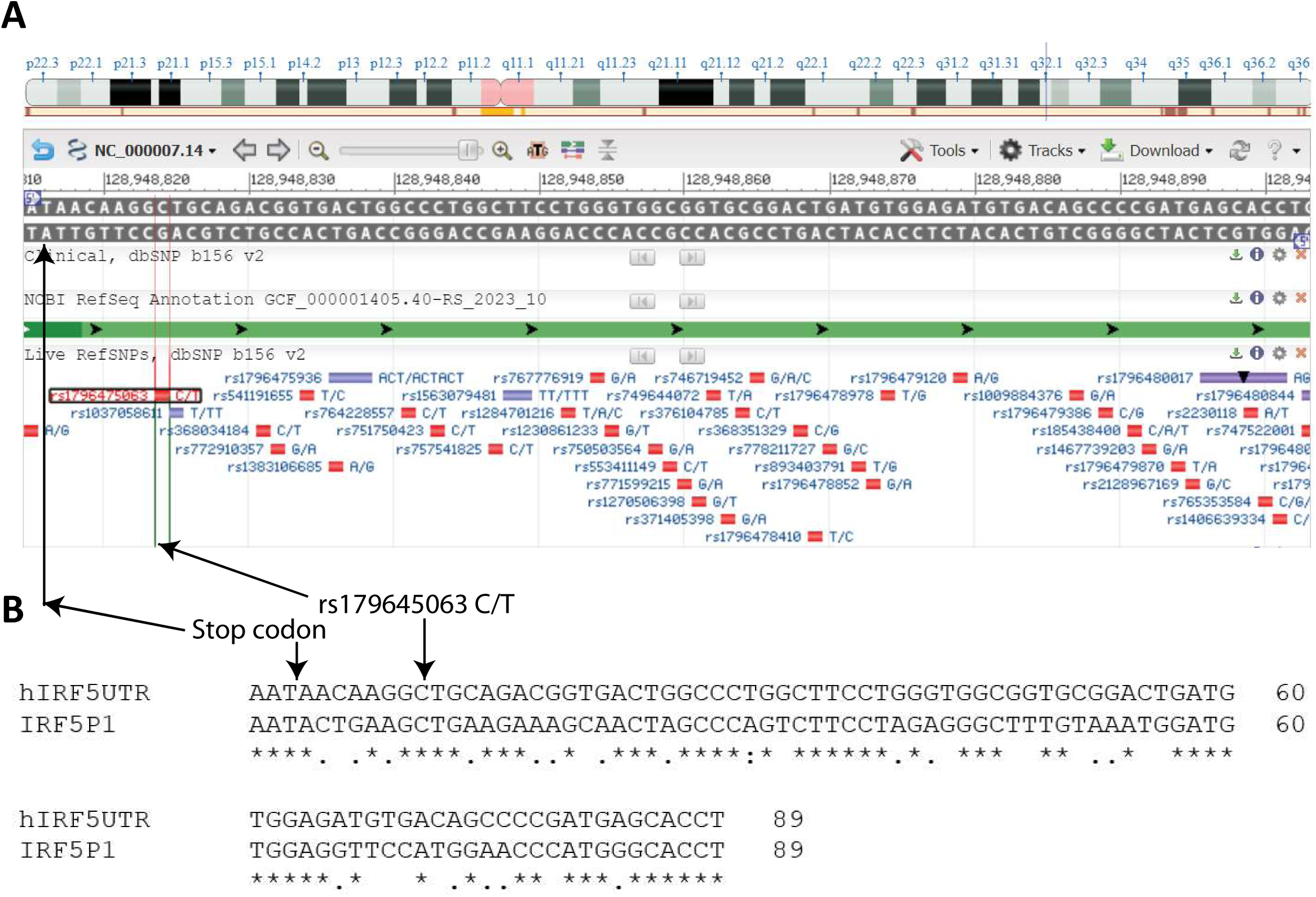
IRF5P1 potentially regulates the IRF5 gene. **Panel A: Variation in the 3’ UTR region of the hIRF5 gene.** The variable sequences, indicated by a red square and corresponding blue alphanumerical identification, represent regions that differ among individuals but do not constitute a mutation. The picture was directly captured in NCBI Variation Viewer at indicated nucleotides. **Panel B: Comparison between the hIRF5 5 UTR and IRF5P1.** The comparison of these sequences was generated using Clustal Omega – Multiple Sequence Alignment. The arrow emerging upward from the labeled stop codon indicates that the hIRF5UTR is identical to the sequence in gray on the ‘A’ figure. The variation changing ‘C’ to ‘T’ renders antisense IRF5P1 unable to bind to hIRF5UTR. Other variable sequences may have similar or opposite effects.

## Discussion

There are indications that primate innate immunity and inflammation pathways have been evolutionary shaped, perhaps in adaptations to primate-specific pathogens, microbiomes, and diets. IRF5 is a key molecule in both inflammation and innate immunity, and comparative studies in IRF5 regulations in primates will likely reveal more about these adaptations.

The primate lineage dates back approximately 65 million years. The presence of a pseudogene only in higher primates suggests that it likely originated and was integrated into primate genomes relatively recently. Our data suggest that IRF5P1 likely originated from retroviral gene transfer from another animal around 60 million years ago (Figure 2). It is interesting that the rhino has a similar gene in its genome, which might be related to another retroviral infection event, but not only kept in the rhinos (Supplemental Figure 1).

The fact that the IRF5P1 pseudogene has been conserved over millions of years of primate evolution implies that it may have some beneficial or functional role specifically in primates. If it were an unimportant nonfunctional sequence, the accumulation of mutations would likely have degraded and removed it from primate genomes over evolutionary timescales.

IRF5P1 could fine-tune IRF5 transcription or translation rates through small RNA regulatory mechanism. It is apparently IRF5P1 antisense are expressed, at least as an intronic RNA of TRIQK (Figure 3). Antisense pseudogene RNA has regulatory effects on parental gene (Hirotsune et al., 2003). In addition, it is well established that introns can serve as sources of regulatory small RNAs like microRNAs (miRNAs) or small interfering RNA (siRNA) in human genes (Mattick & Makunin, 2005; Rearick et al., 2011). Therefore, it is quite possible that antisense IRF5P1 RNA expression offers a novel source to generate some small RNAs. Those RNAs will most likely target IRF5 for fine tuning or other regulations (Figure 4). Of note that miRNAs are well established regulators for IRF5 (Chang et al., 2021; Chen et al., 2017; Fang et al., 2021; Gong, Guo, & Zhang, 2018; Lin et al., 2022; Wu et al., 2021). IRF5P1 may contribute to the optimization of small RNA-mediated immune modulation driven by selection pressures from primate-specific pathogens.

Although sense IRF5P1 has limited expression in healthy individuals (Supplemental Figure 4), it is possible that IRF5P1 may be expressed in a pathogenic situation. In addition, despite its pseudogene classification, IRF5P1 could potentially be gaining primate-restricted open reading frames and expressing truncated or altered peptide sequences that carry out primate-specific functions. Because IRF5 is a critical component of inflammation and innate immunity, regulatory effects of IRF5P1 on components of innate or adaptive immunity could help tailor immune system functionality to primate-specific pathogens. Further molecular investigations of IRF5P1 expression patterns, interactions, and functionality would help elucidate the significance this intriguing primate pseudogene acquired since its emergence. Understanding the role of regulatory pseudogenes in the functional landscape of interferon and inflammatory signaling will offer novel insights into transcriptional control mechanisms and the pathogenesis of related diseases.

## Materials and Methods

### Searching of IRF5P1 genes

The DNA sequences of the human IRF5P1 or other relevant mRNA and protein sequences were used as the queries (see Supplementary Tables 1 for accession numbers) to perform nucleotide or protein similarity searches using BLASTN (nucleotides) and BLASTP (protein) against the non-redundant protein database at NCBI with the default options. Sequences were collected. When more than one isoforms were available from the same species, one isoform that was most similar to those from other species was selected. Protein sequences that were partial and too short were excluded.

In addition, UCSC Genome Browser BLAT Search was used for searching for genomic sequence similar to IRF5P1 (https://genome.ucsc.edu/index.html).

### Phylogenetic analysis of nucleotides and proteins

The nucleotide sequences were aligned using MAFFT v7 with the E-INS-i iterative refinement method. The alignments were downloaded with festa format and used for the maximum likelihood phylogeny was reconstructed using IQ-Tree release 1.6.11 with the default options and the amino acid substitution model estimated. Brach support values were obtained using the ultrafast bootstrap support and SH-aLRT branch test both with 1000 replicates. The final alignment was performed using each specific alignment as the profile and using the “merge” alignment of MAFFT with the E-INS-i iterative refinement method. The visualization of the phylogenies was performed using Interactive tree of Life (ITOL) website (https://itol.embl.de/) (Letunic & Bork, 2021).

---

**Supplemental Table 1:**
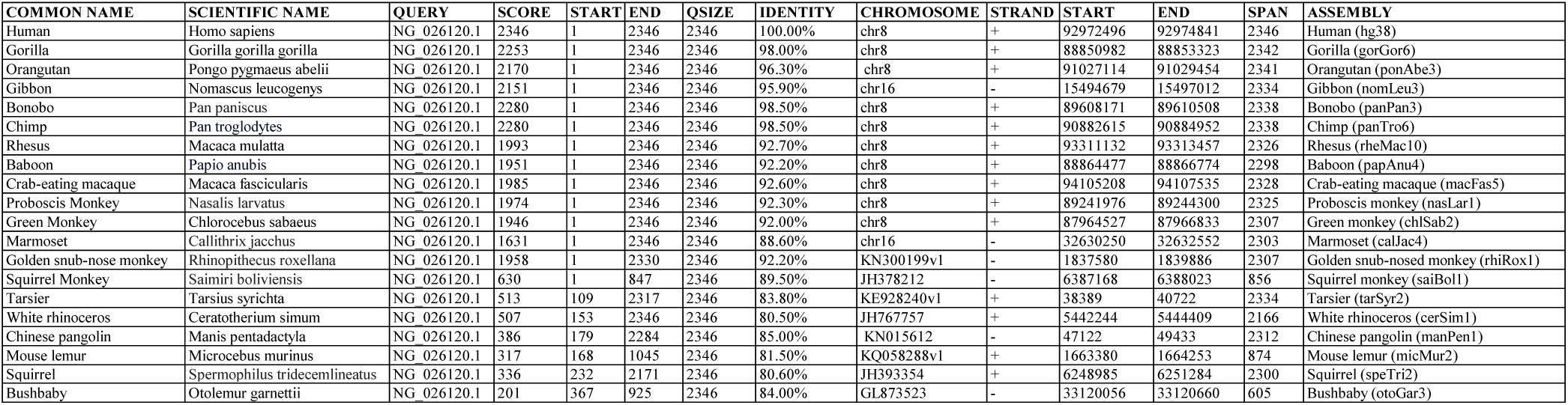
The genomic sequences used for this study.

**Supplemental Figure 1:**
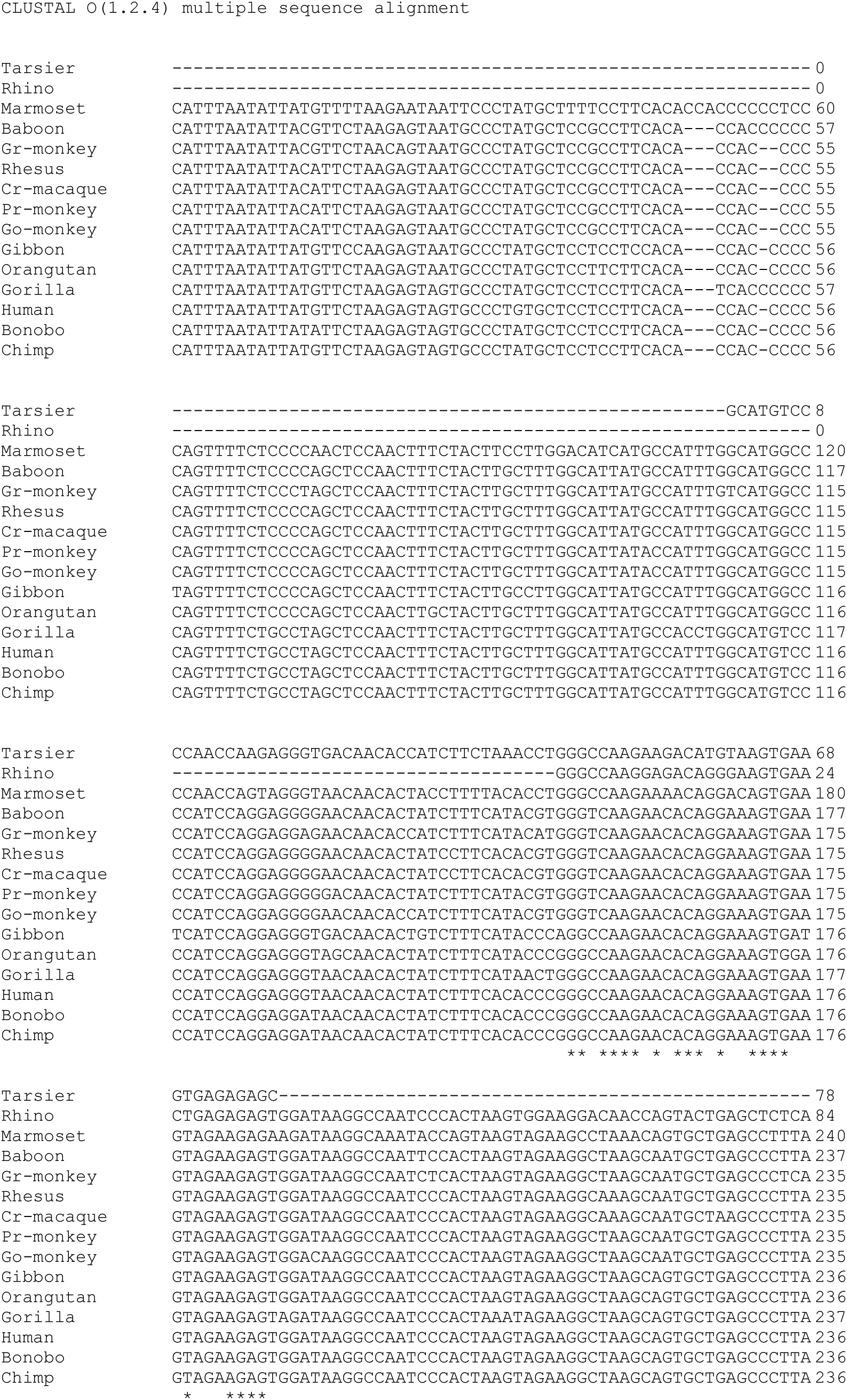
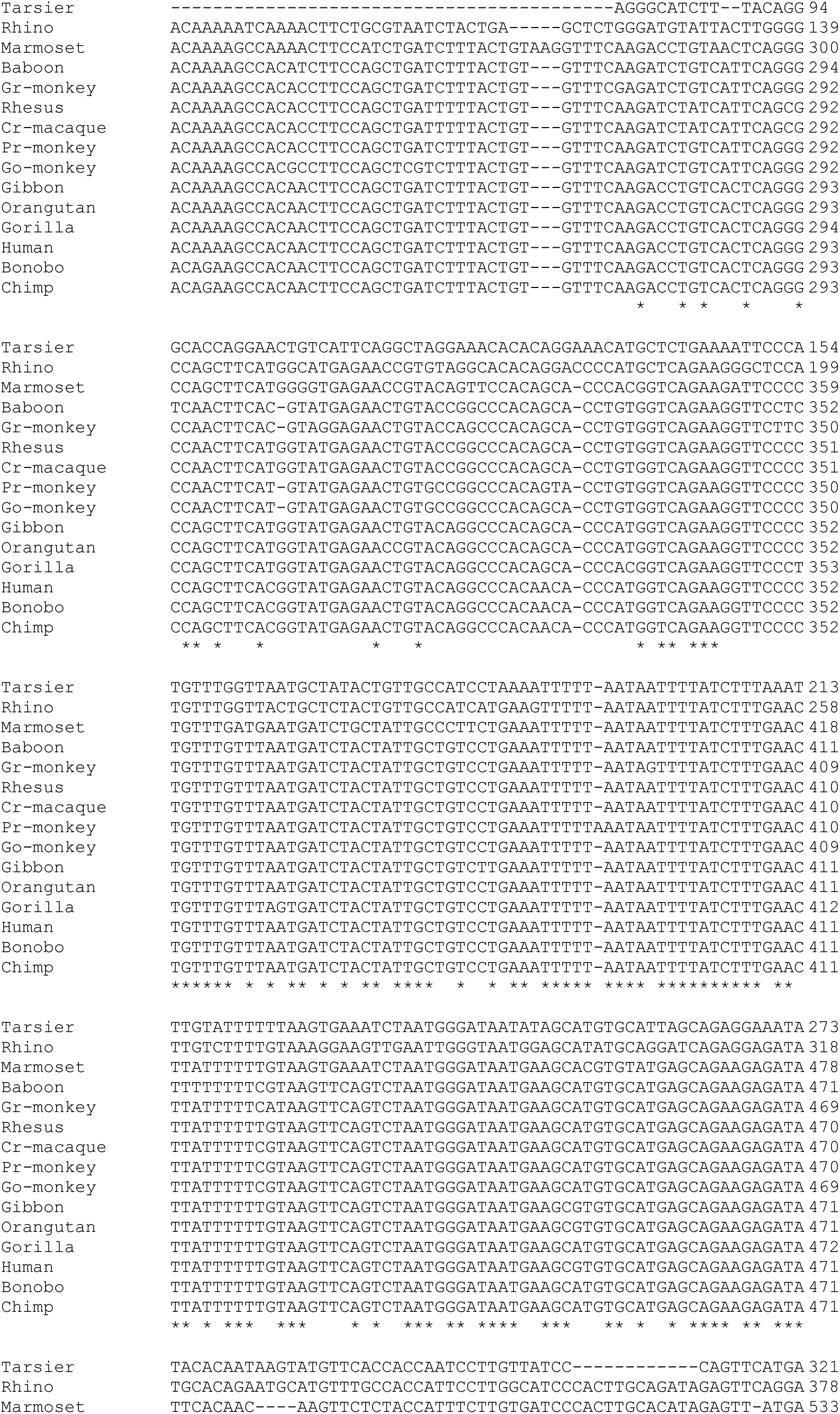
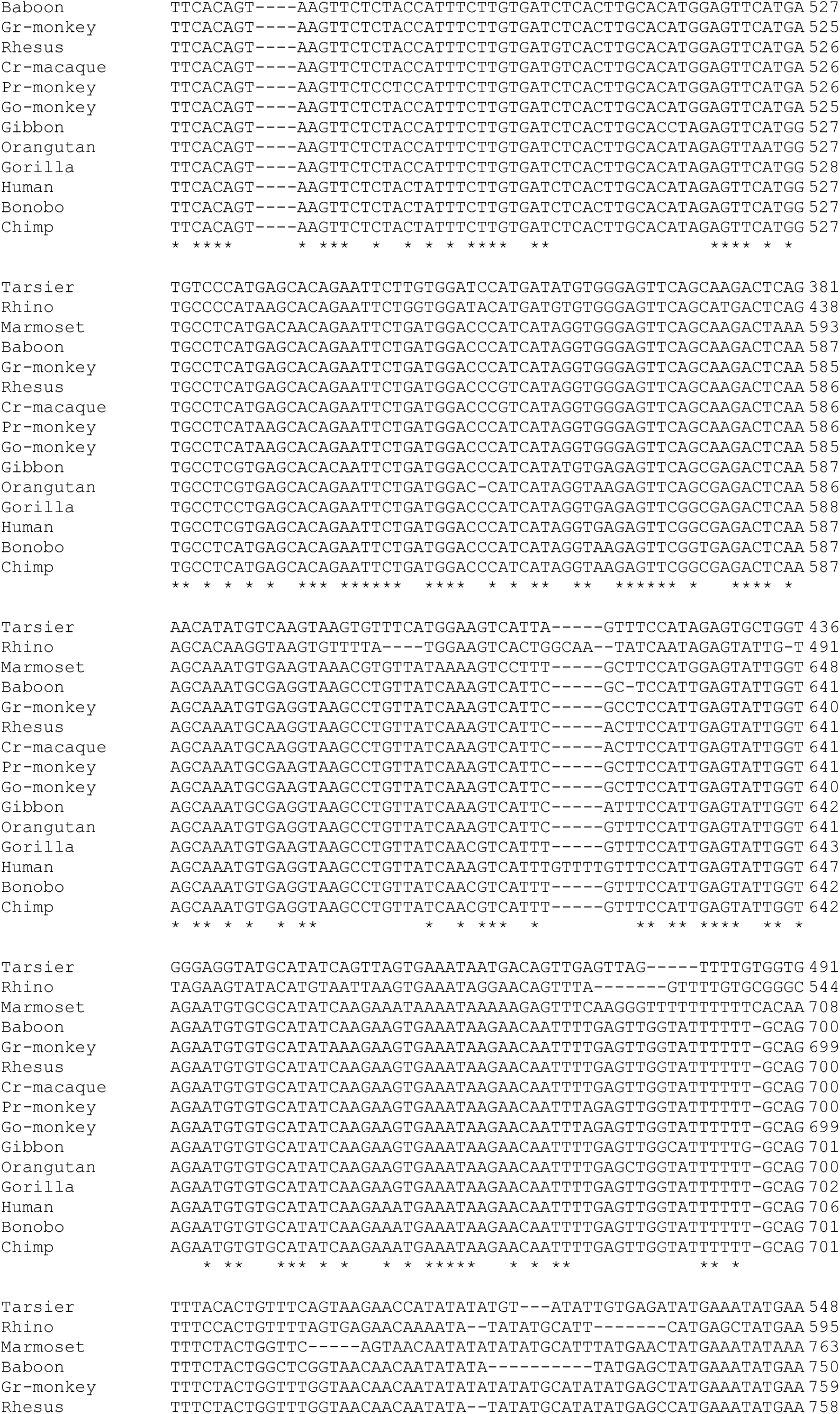
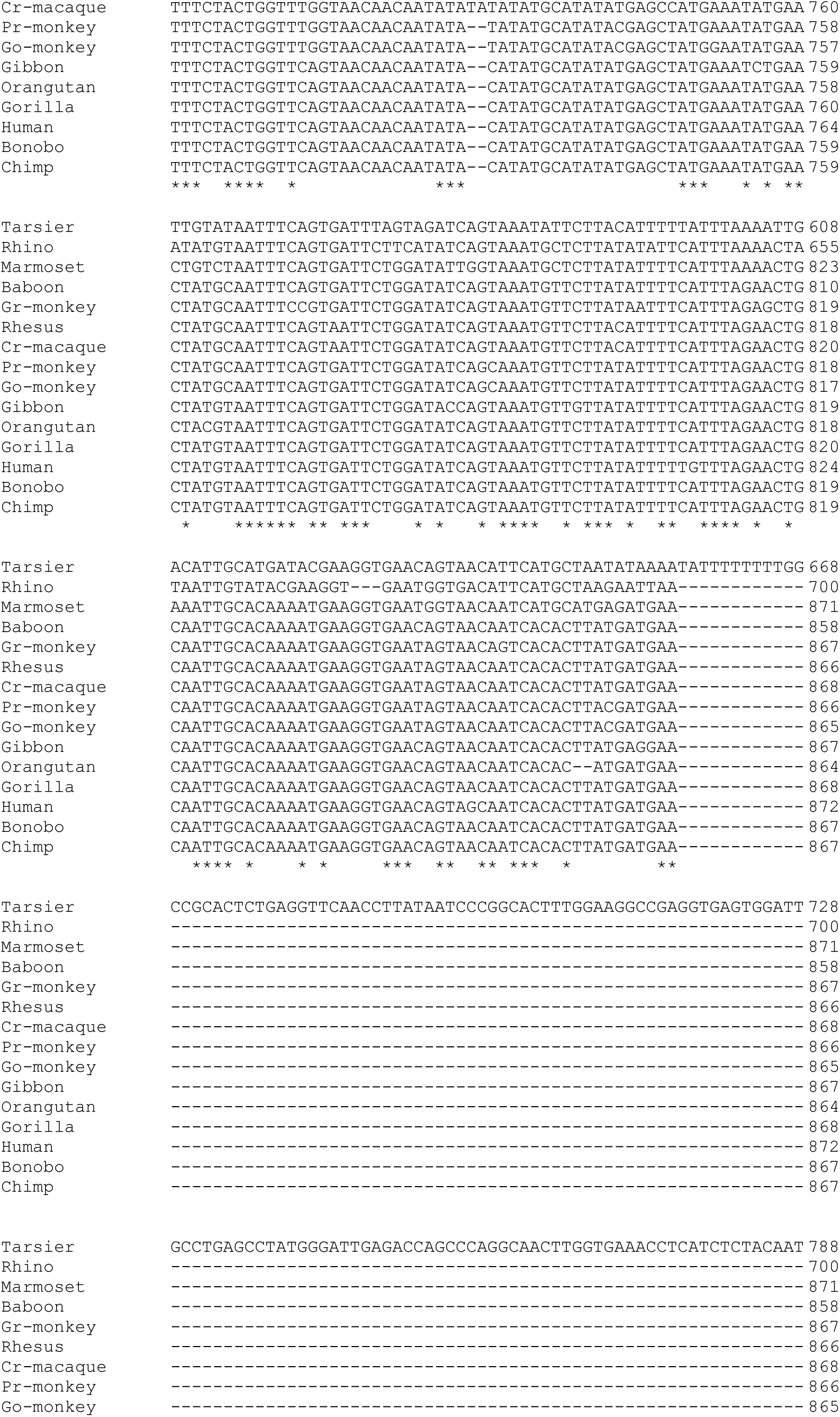
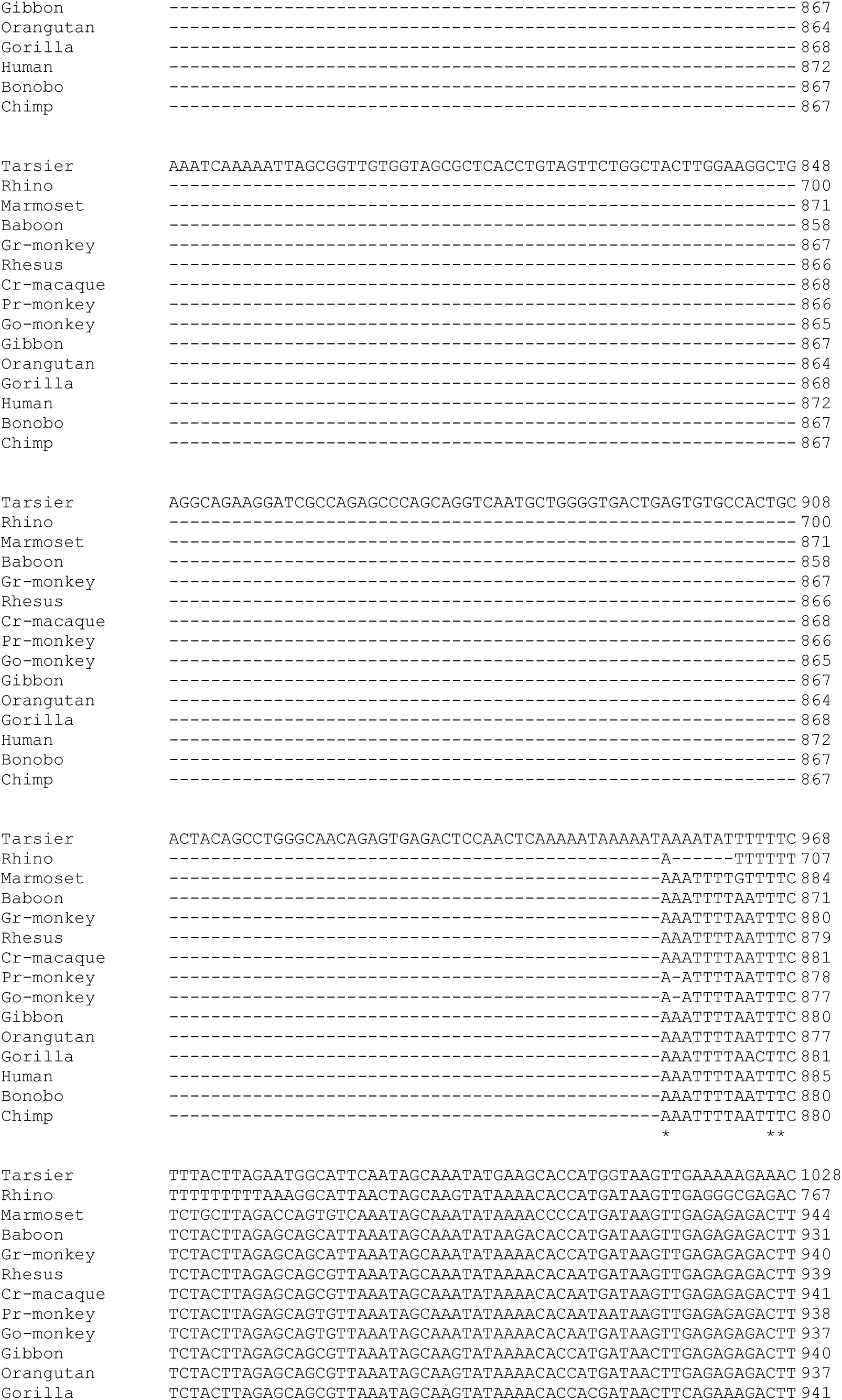
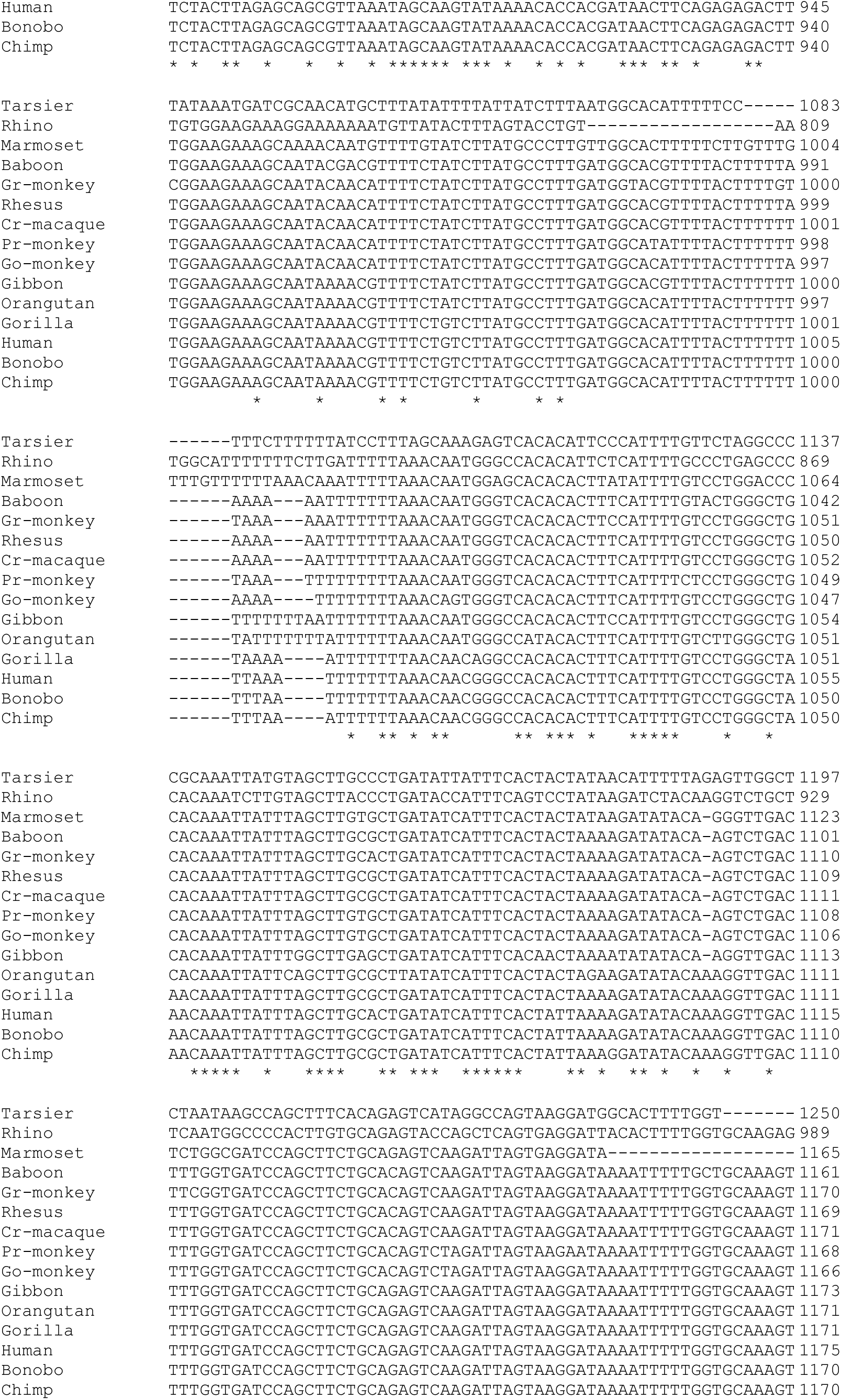
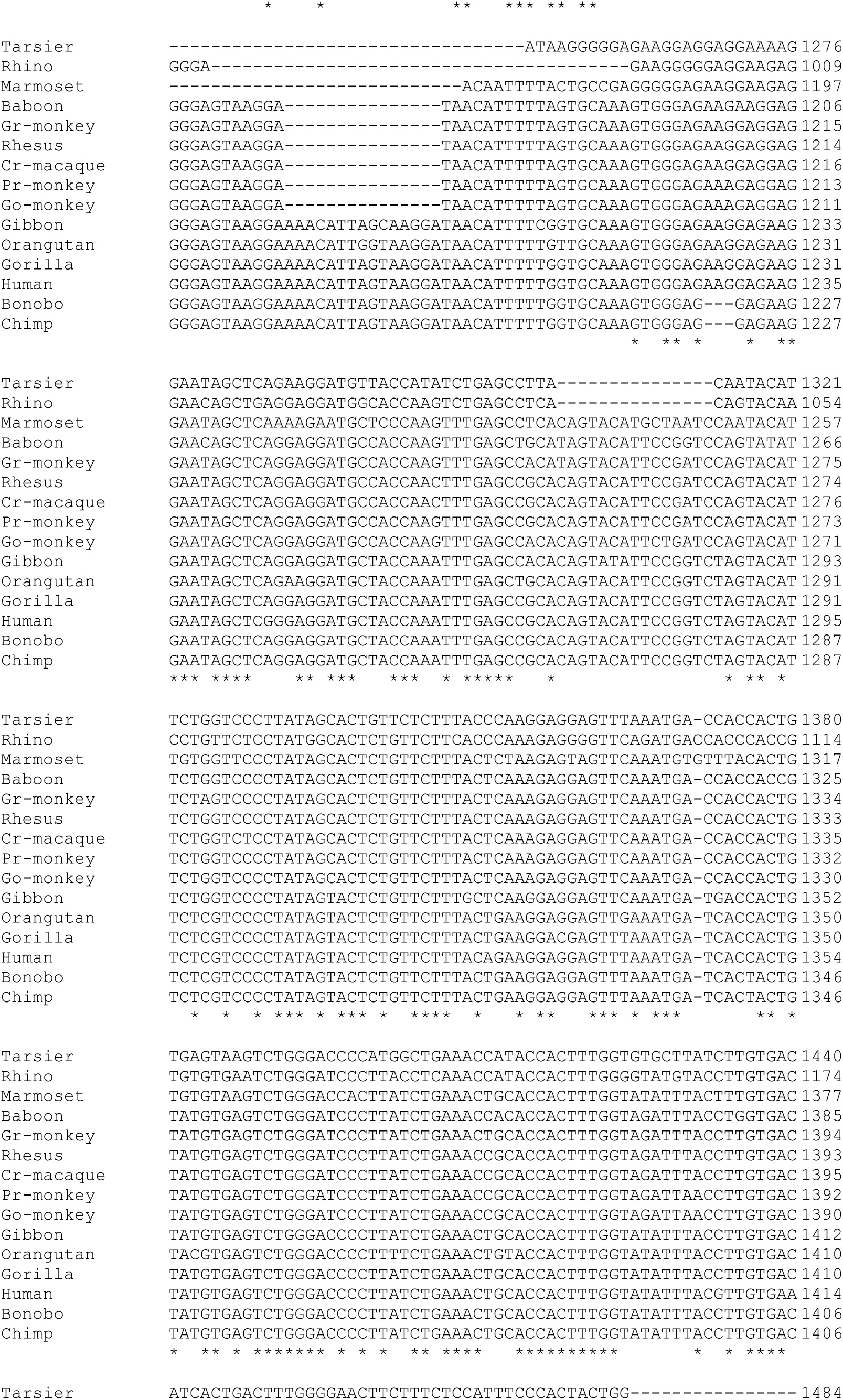
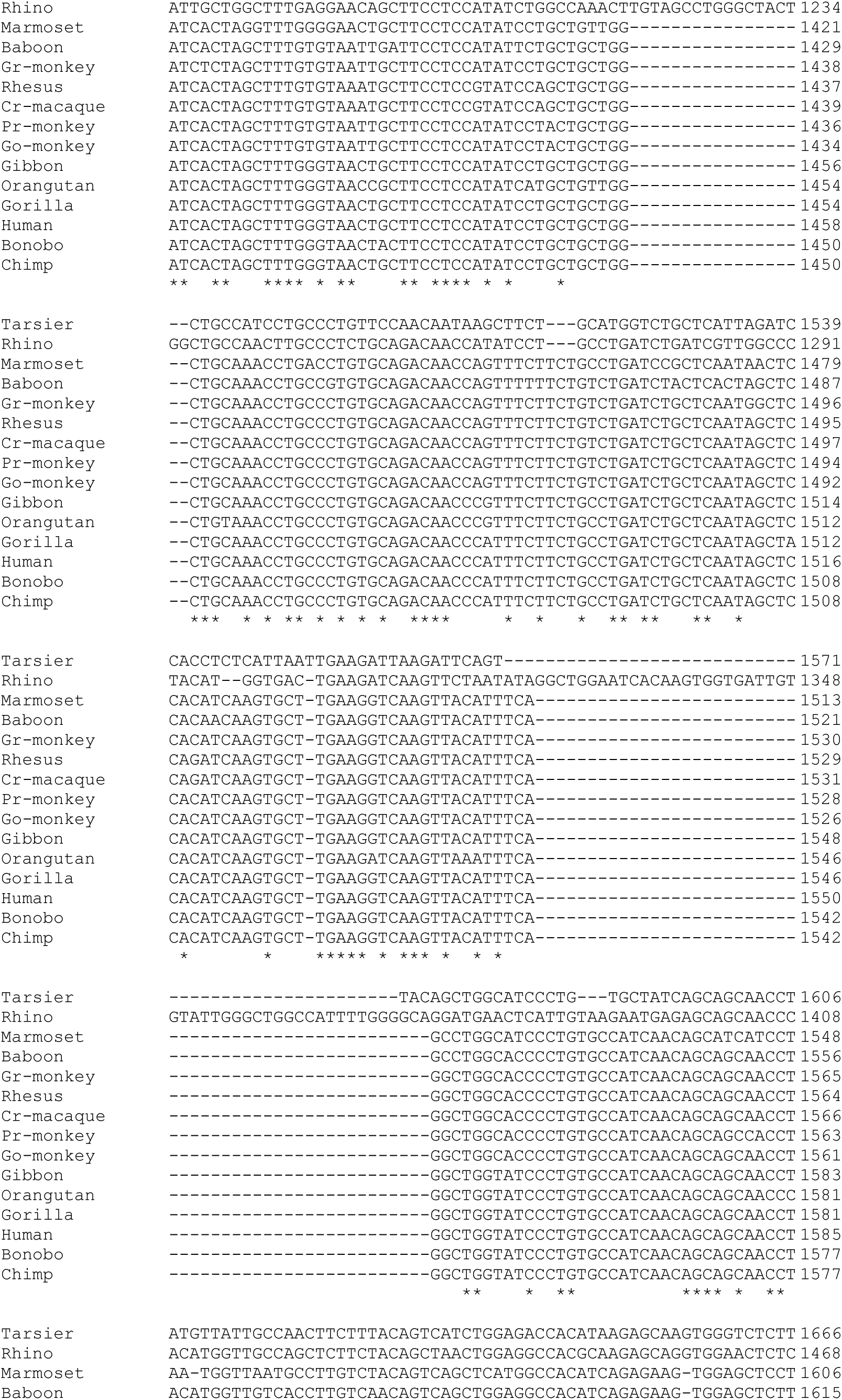
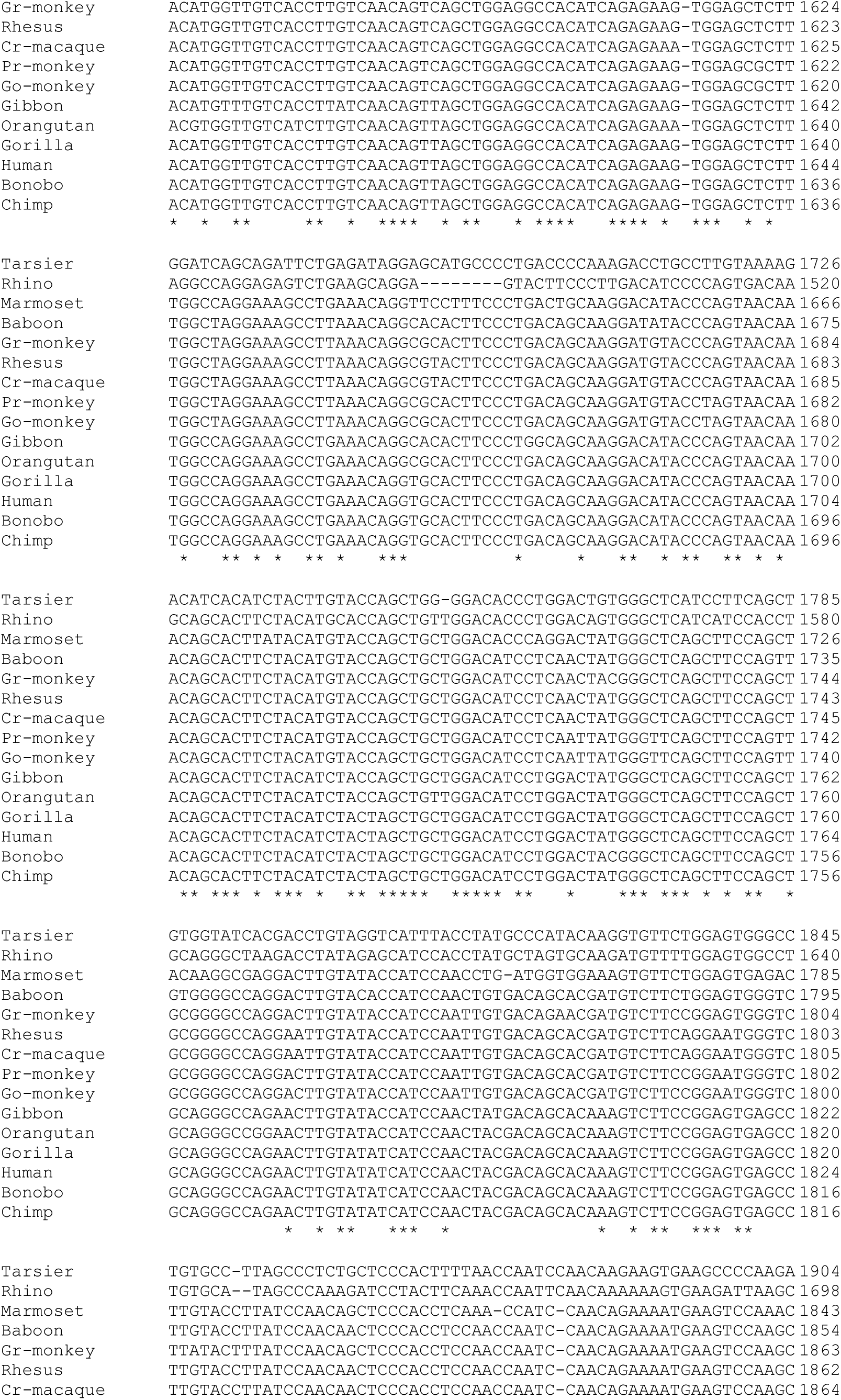
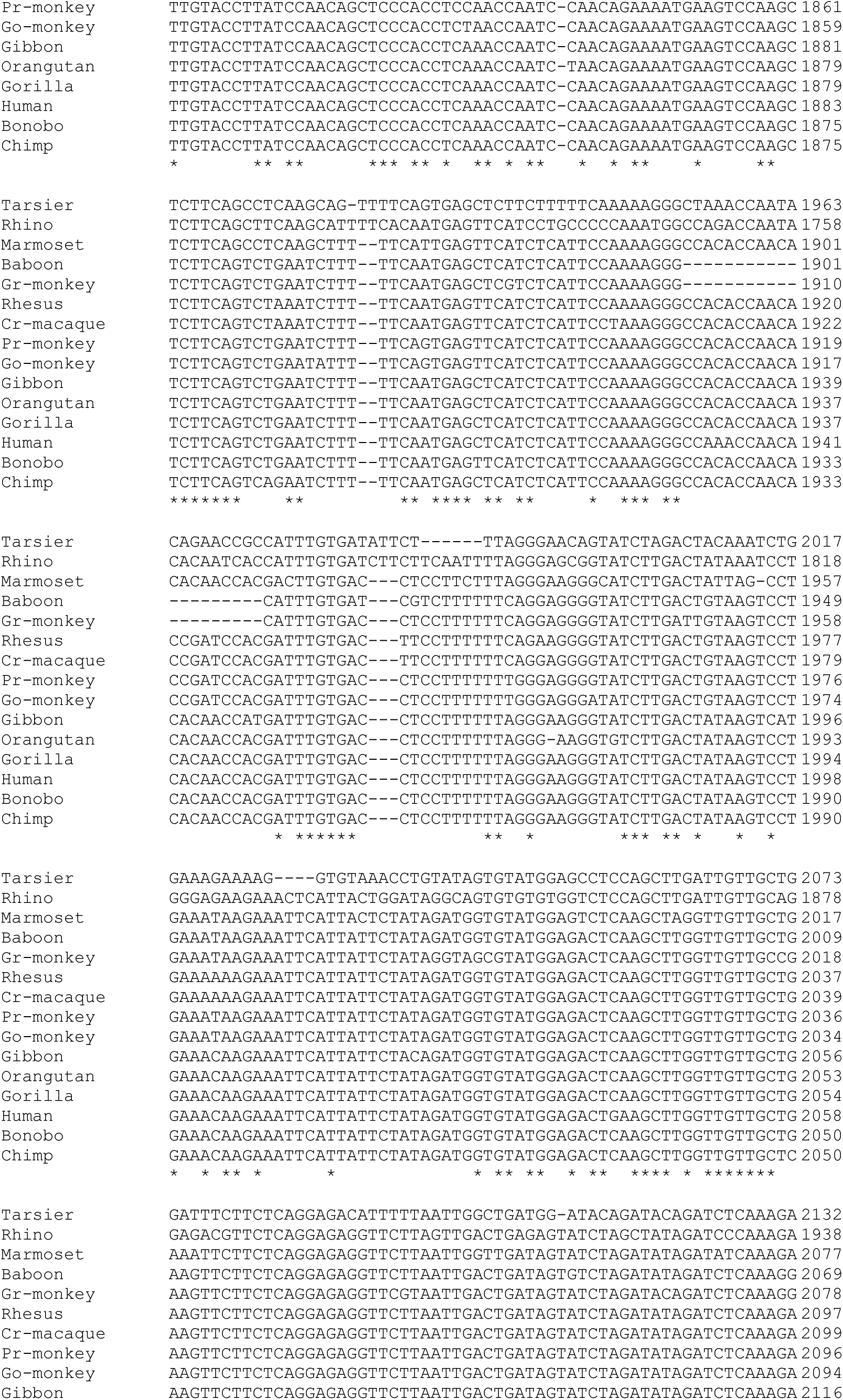
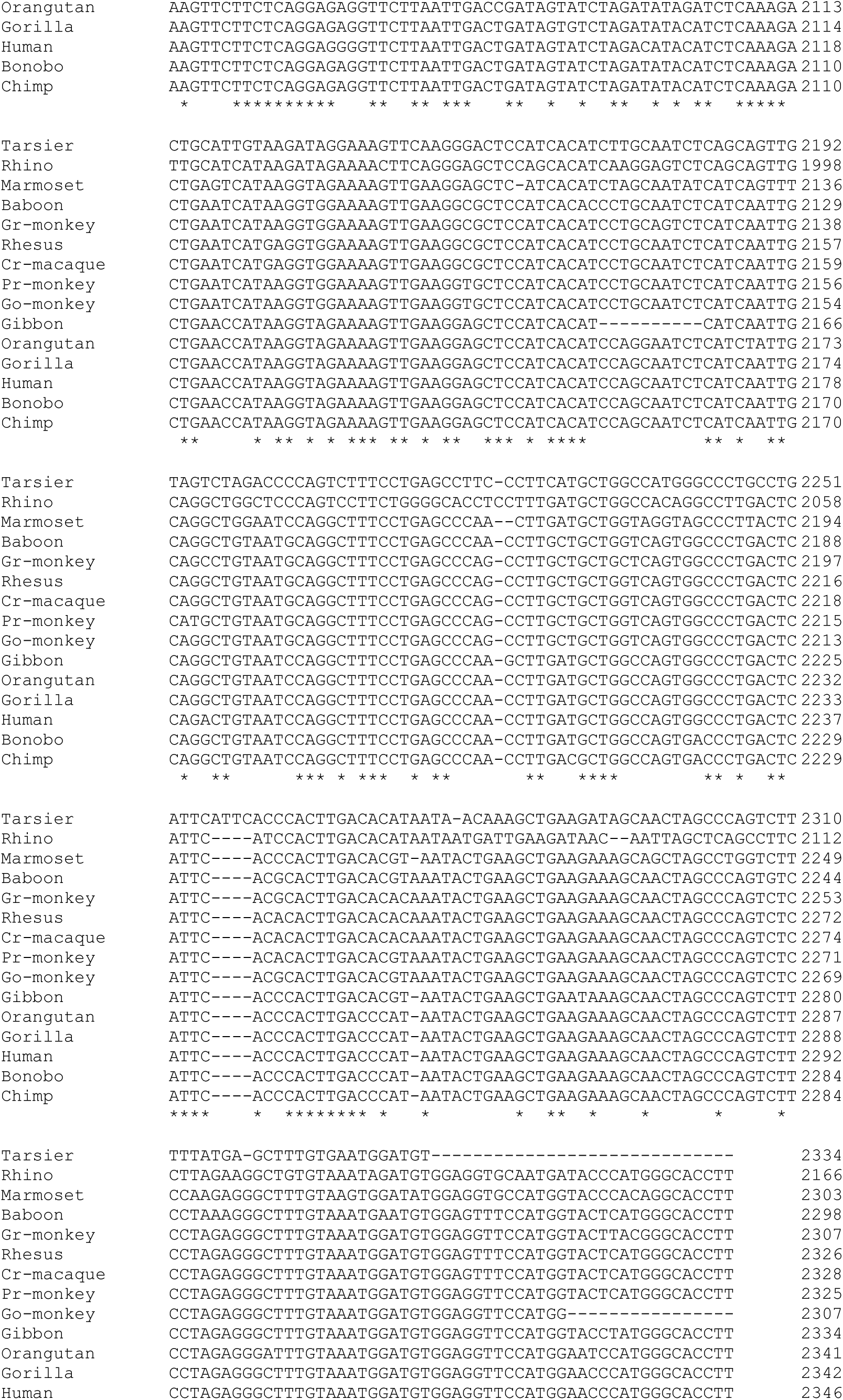
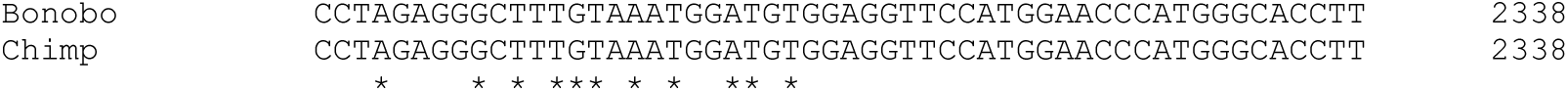
Clustal Omega multiple sequence alignment of the 15 sequences in Supplemental Table 1. Abbreviations for species names: Cr-macaque: crab-eating macaque; Go-monkey: golden snub-nose monkey; Gr-monkey: green monkey; Pr-monkey: proboscis monkey.

**Supplemental Figure 2:**
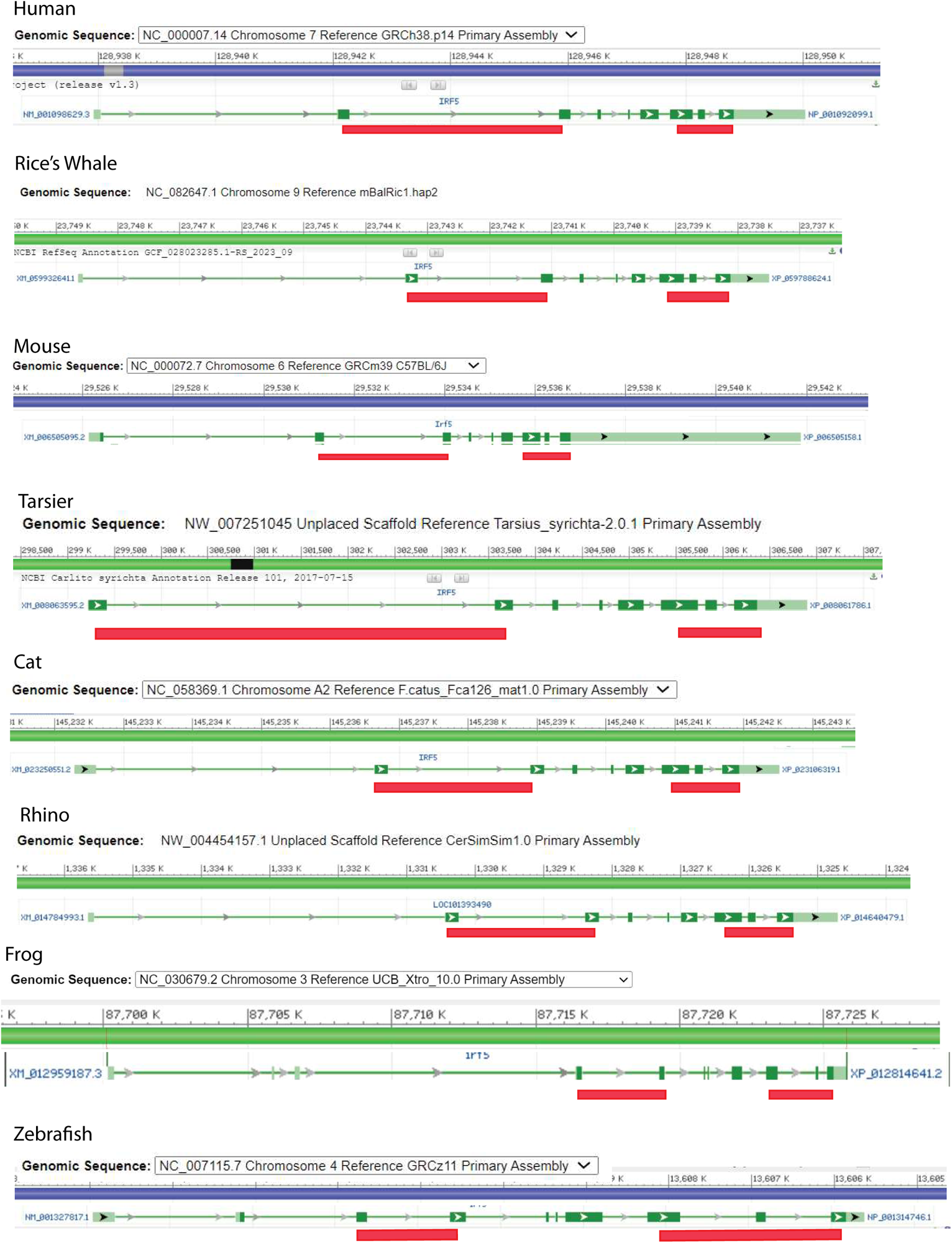
Gene Structures and Transcripts of IRF5 in Various Species. The common species name is listed at the top of the diagram. The source of genomic sequences, chromosome numbers, and nucleotide coordinates is shown. Light green bars represent exon sequences, and solid green bars represent protein-coding sequences. The left red line represents the corresponding sequences for Section 2, while the right red line represents the corresponding sequences for Section 4. Section 2 is a combination of two exons, while Section 4 is a combination of three exons. The diagrams are directly obtained from the NCBI gene report pages. The scales of the diagrams have been adjusted for comparison purposes among species.

**Supplemental Figure 3:**
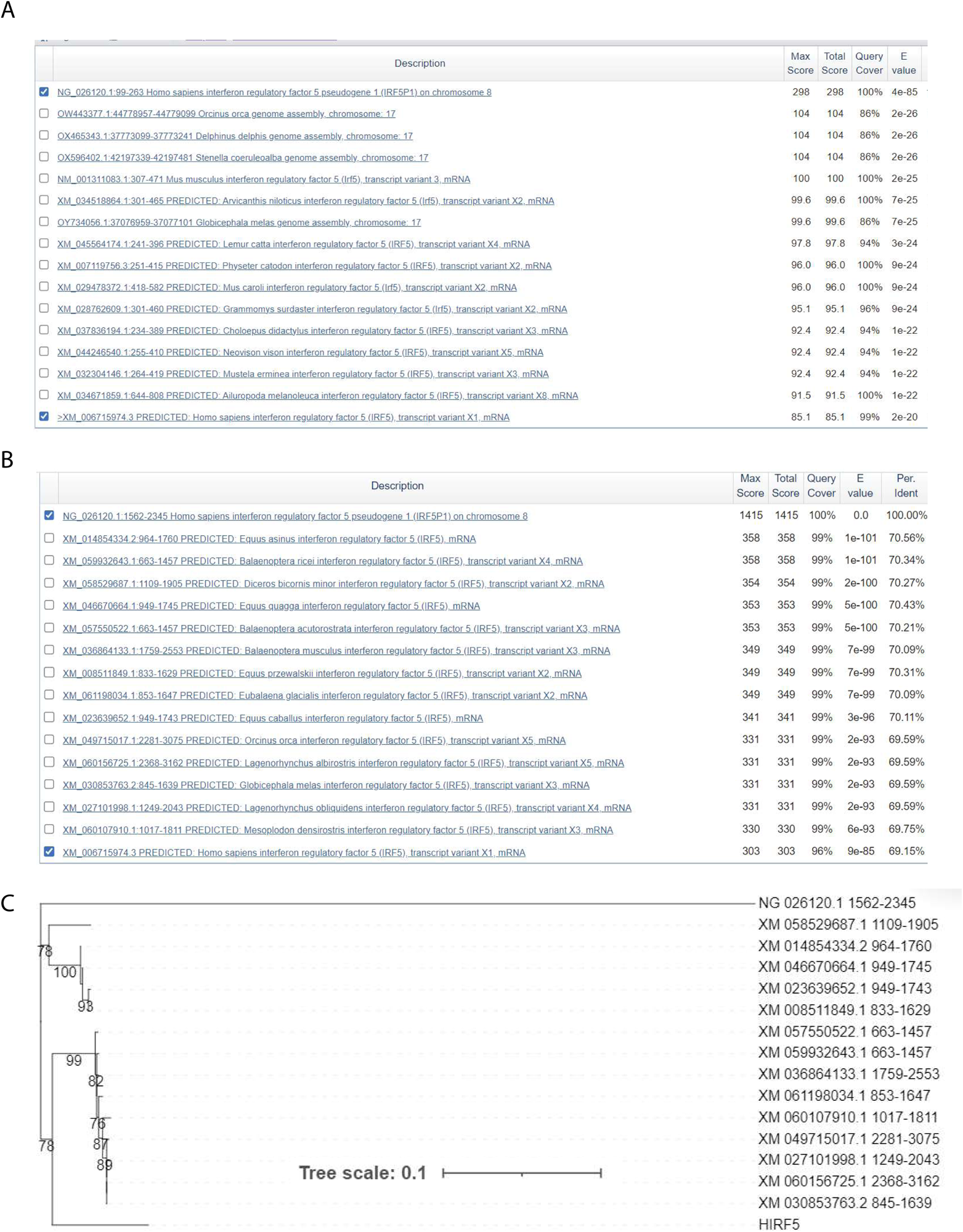
**Panel A**: A BLAST was generated utilizing Section 2 (99-263) of the human IRF5P1. Query cover, E value, and percent identity values are displayed. The human IRF5 sequence is highlighted at the bottom of the blast. **Panel B**: Same as Panel A but the input sequence was Section 4 (1562-2345) of the IRF5P1. **Panel C**: Phylogenetic relationships among IRF5P1 and IRF5 mRNA sequences in various species. The maximum likelihood phylogeny was reconstructed using IRF5P1 nucleotide (Section 4). Internal nodes supported by 70% or higher bootstrap values are indicated by numerical values on nodes.

**Supplemental Figure 4:**
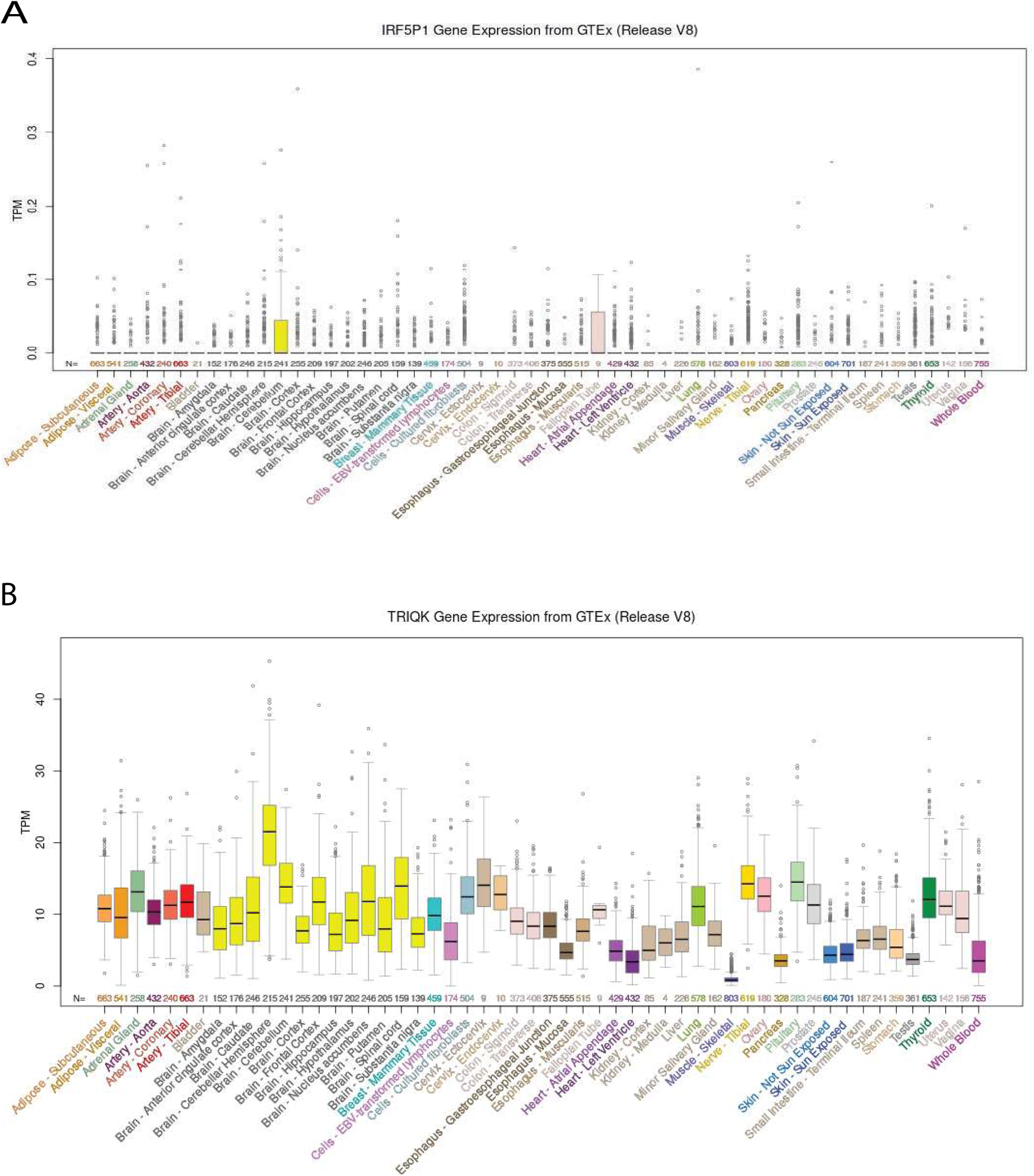
Gene Expression from various cells and tissues. Gene expression levels are measured by transcripts per million (TPM). **Panel A**: IRF5P1; **Panel B**: TRIQK Gene expression. The results are based on hg38 Gene Expression in 54 tissues from GTEx RNA-seq of 17382 samples, 948 donors (V8, Aug 2019) (Lonsdale et al., 2013). Ensembl gene ID: ENSG00000251347.2 (IRF5P1) and ENSG00000205133.11(TRIQK).

## Notes

### Competing Interest Statement

The authors have declared no competing interest.

